# The axonal sorting activity of pseudorabies virus Us9 protein depends on the state of neuronal maturation

**DOI:** 10.1101/2020.08.06.239384

**Authors:** Nikhila S Tanneti, Joel D Federspiel, Ileana M Cristea, Lynn W Enquist

## Abstract

Alpha-herpesviruses establish a life-long infection in the nervous system of the affected host; while this infection is restricted to peripheral neurons in a healthy host, the reactivated virus can spread within the neuronal circuitry, such as to the brain, in compromised individuals and lead to adverse health outcomes. Pseudorabies virus (PRV), an alpha-herpesvirus, requires the viral protein Us9 to sort virus particles into axons and facilitate neuronal spread. Us9 sorts virus particles by mediating the interaction of virus particles with neuronal transport machinery. Here, we report that Us9-mediated regulation of axonal sorting also depends on the state of neuronal maturation. Specifically, the development of dendrites and axons is accompanied with proteomic changes that influence neuronal processes. Immature superior cervical ganglionic neurons (SCGs) have rudimentary neurites that lack markers of mature axons. Immature SCGs can be infected by PRV, but they show markedly reduced Us9-dependent regulation of sorting, and increased Us9-independent transport of particles into neurites. Mature SCGs have relatively higher abundances of proteins characteristic of vesicle-transport machinery. We also identify Us9-associated neuronal proteins that can contribute to axonal sorting and subsequent anterograde spread of virus particles in axons. We show that SMPD4/nsMase3, a sphingomyelinase abundant in lipid-rafts, associates with Us9 and is a negative regulator of PRV sorting into axons and neuronal spread, a potential antiviral function.

**Author Summary:** Viral pathogenesis often is age-dependent, with more severe outcomes for infected fetuses and neonates compared to adults. As neurons age and mature, dendrites and axons polarize with distinct functions that affect neurotropic virus replication and neuronal spread of infection. This study investigates how neuronal maturation of peripheral nervous system neurons, the site of alpha-herpesvirus life-long latency and reactivation, affects replication and neuronal spread of pseudorabies virus. Characterization of infected immature and mature primary cultures of superior cervical ganglionic neurons revealed significant differences in protein composition and cellular processes that affected the activity of Us9, a viral protein required for sorting virus particles into axons. We identified neuronal and viral proteins that interact with Us9 in immature and mature neurons. Among these, we demonstrate that SMPD4/nsMase3, a sphingomyelinase critical for membrane organization and neuronal function, regulates PRV neuronal spread by preventing capsid association with Us9-containing membranes, presenting a possible antiviral function.

## Introduction

Pathogenesis of neurotropic virus infections is influenced by age of the infected host. For example, the recent epidemic of Zika and West Nile virus infections highlighted the extreme disease outcomes in infected fetuses such as microcephaly and mental retardation, in comparison to less pathogenic infections in the adult population (1, 2). Similarly, pathogenic outcomes of alpha-herpesvirus infections, such as those caused by herpes simplex virus, varicella-zoster Virus and pseudorabies virus also are age-dependent. In these cases, infections in newborns are more severe than infections of adults (1, 3, 4). This study explores the relationship between neuronal age and the clinical outcome of alpha-herpesvirus infection.

Pseudorabies virus (PRV), like other alpha-herpesviruses, establishes a life-long, latent infection in the peripheral neurons of the host. Stress of many types can reactivate the quiescent viral genome, prompting the assembly of new virus particles that move from neuronal cell bodies to axons that innervate peripheral sites where infection of epithelial tissues promotes the spread of infection to uninfected hosts. In adults, this unique process of spread occurs in a fully mature nervous system. In immature hosts, such as newborns with still developing nervous system, neuronal mechanisms that restrict virus spread may not be developed. As a result, the unregulated spread of the virus throughout the neuronal circuitry, including the central nervous system, could lead to encephalitis or systemic spread. The property of alpha-herpes virus infection neuronal-spread being exploited by neuroscientists to map the neuronal-circuity and by oncologists to treat brain tumors (5, 6). However, the underlying mechanisms of how the virus undergoes neuronal spread in mature or immature neurons are not well understood.

Anterograde spread of infection in a mature neuron comprises at least two different processes. The first step, called axonal sorting, is a highly regulated process in which only a small fraction of mature virus particles that assemble in the cell-body move into the axon. In the second step, called anterograde transport, the virus particles are transported on microtubules by kinesin motors down the axon to sites of particle egress. The two steps are necessary for successful spread of virus from the infected neuron to connected cells. (7)

Axonal sorting of PRV particles into axons requires the viral protein Us9, a type-II transmembrane protein that is present in neuronal ER and Golgi membranes as well as in the viral envelope during infection. As the topology of type-II membranes suggests, the N-terminus 68 amino acids of Us9 are exposed to the cytoplasm where Us9 contacts neuronal proteins (8). Kramer *et al* showed that Us9 interacts with Kif1a, a microtubule plus-end directed motor (9) to mediate virus transport. This mechanism is conserved among alpha-herpesviruses and is strengthened by other Us9-associated viral proteins such as gE and gI (10); however, the roles of other neuronal proteins in this mechanism are less understood. Importantly, the Us9 protein must be located in detergent-resistant-membranes or lipid-rafts to interact with Kif1a and move virus particles into axons (11). Lipid-rafts are important for the function of neurons as well as pathogenesis of many viruses that utilize cellular membranes (12).

Here, we compared infections of PRV with wildtype and mutant Us9 protein to identify virus-host interactions underlying axonal sorting, anterograde transport, and neuronal spread of infection from axons to epithelial cells. We developed primary cultures of superior cervical ganglia (SCG) representing immature and mature stages to understand how neuronal development affects Us9 function. We also employed mass-spectrometry techniques to identify interactions between PRV-Us9 and host proteins in immature and mature neurons. We determined that the Us9 interacting host protein, SMPD4, has an anti-viral function by negatively regulating anterograde spread of PRV in axons.

## Results

### Establishing Mature and Immature Cultures in vitro

To understand how neuronal age affects PRV biology, we established and characterized primary cultures of immature and mature neurons from superior cervical ganglia (SCG), sympathetic neurons of the peripheral nervous system that are naturally infected in the affected hosts (8, 13, 14). SCGs from 17.5-day embryonic rats were harvested and cultured with neuronal growth factor for 3-4 DIV (days in vitro) or 20+ DIV to represent immature and mature neurons, respectively. Multiple characteristics were assayed to determine both the maturity and uniformity of the cultures. Phase-contrast microscopy **(Figure 1A)** comparing the two developmental stages revealed that: (1) neuronal soma/cell-bodies grow in size with maturation, (2) cell-bodies cluster into groups and stop dividing as terminal differentiation occurs, and (3) a robust network of axons forms and expands over time. Immature neurons have modest projections termed neurites that are morphologically distinct from mature axons. Mature cultures form an axonal network that is necessary to establish synaptic connections and produce action potentials (15). The increase in cell body size and the extensive growth of axons were correlated with an increase in protein mass per cell during development **(Figure 1B)**. Immature SCGs at 3 DIV contained an average of 2.5 μg of protein per SCG compared to mature neurons at 21 DIV with an average of 19.2 μg of protein per SCG. It is important to understand that the maturity of neurons is defined arbitrarily by DIV and that induction of maturity is not synchronous. It is likely that immature neurons are a diverse mixture representing neurons in different stages of maturation. However, as neurons age in culture, they become substantially more uniform as they terminally differentiate.

**Figure 1 Legend:**
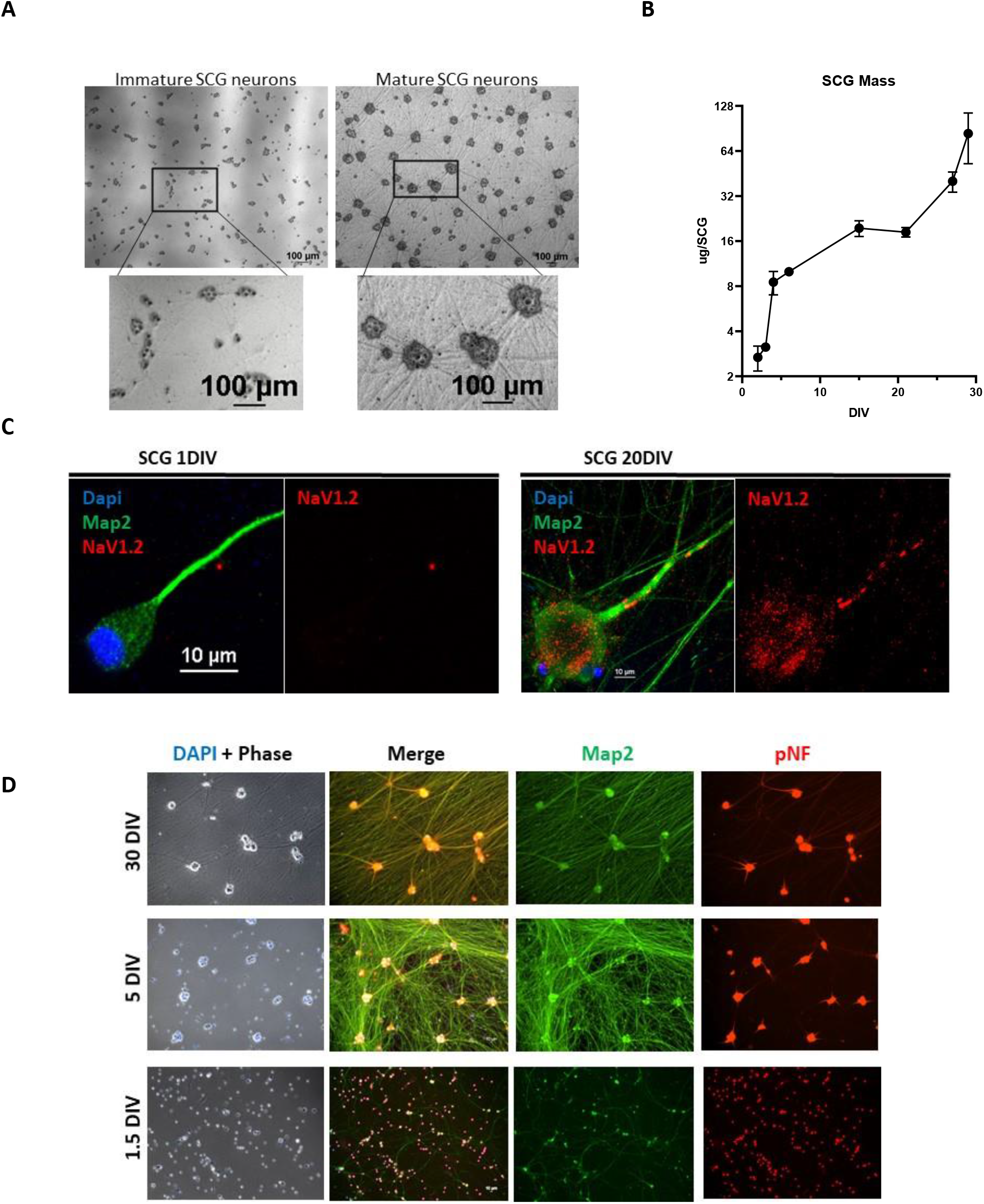
Neuronal Maturation is established by 5 DIV. **A**: Maturation of SCGs is characterized by building a neuronal-network. Phase contrast image comparing immature SCGs to mature SCGs that develop dense axon bundles (straight lines) and clusters of soma (dark grey). **B**: Mass per dissociated SCG at various DIV (days in vitro). Bars represent standard deviation. **C**: Mature neurons express NaV1.2 Immunofluorescence staining in axon. Immature (1 DIV) SCG does not localize NaV1.2 compared to mature (30 DIV) axon localizing Nav1.2. Map2 serves as a neuronal marker and Dapi for nucleus. **D**: SCG maturation is acquired by 5 DIV. Maturation marker pNF (in red) is localized only to the soma (round cell bodies) at 1 DIV, after which localization spreads to the axons (green lines) by 5 DIV, and maintains axonal expression at 28 DIV. Map2, a somato-dendritic marker (in green) is localized to all regions of the axon and does not change with neuronal age.

Neuronal development is accompanied by terminal differentiation and specialization of distinct functions, such as the polarization of axons and generation of action potentials (16). We examined two proteins that serve as markers of maturation, NaV1.2 -a sodium channel necessary for the generation and propagation of action potentials, and the axonal marker phospho-Neurofilament H, pNfH (17, 18). Immunofluorescence microscopy experiments confirmed that the late maturation maker, NaV1.2, localizes to the proximal axon by 20 DIV **(Figure 1C)**, suggesting that maturation is completed by this time. To understand when maturation takes place, we looked at pNfH localization over-time **(Figure 1D)**. While pNfH is localized only to the cell-bodies in the 1.5 DIV SCG neurons, the localization expanded into the proximal axonal regions around 5 DIV and by 30 DIV pNfH is localized throughout all axons. This observation suggests that neuronal maturation is likely complete around 5 DIV, which is earlier than previously thought. Together, these results define the time-frame of SCG neuronal maturation allowing the study of virus infection in the context of neuronal age.

### Efficient Axonal-Sorting of Pseudorabies Virus Particles Requires Neuronal Maturation

Previously, Tomishima and Enquist [17] found that that neuronal maturity affected Us9-dependent sorting of viral structural glycoproteins into SCG axons. We hypothesize that neuronal maturity also affected the axonal sorting of mature PRV virions into SCG axons. To visualize virus particles, we constructed dual color PRV recombinants expressing an mRFP-VP26 fusion protein and a GFP-Us9^WT^ fusion protein or the mutant GFP-Us9^YY^ fusion protein, a non-functional missense protein (9, 19). This mutant Us9 protein contains a di-tyrosine to di-alanine substitution, which does not affect Us9 protein expression or lipid-raft localization, but the missense protein does not interact with Kif1a and fails to sort virus particles into axons (20). Thus, the comparison of the two infections will identify Us9-associated proteins specific to the function of axonal sorting and anterograde neuronal spread of virus.

Immature and mature dissociated SCG neurons were infected at high MOI with the dual color recombinant PRV mutants expressing mRFP-VP26 and either GFP- Us9^WT^ or GFP-Us9^YY^. At 12 hours post-infection (hpi), the number of viral particles visible as red mRFP-VP26 (capsid protein) puncta in the proximal segment of axons were measured, as assessed by mRFP-VP26 fluorescence **(Figure 2A)**. Infections of mature SCG neurons with recombinant PRV expressing Us9^WT^ resulted in sorting of an average of 30 virus particles into axons, in comparison to 2 particles found in axons after infection with the recombinant expressing mutant PRV Us9^YY^ **(Figure 2B)**. The observed >90% reduction in sorting is consistent with previous findings that Us9 is required for axonal sorting of PRV particles in mature axons (20).

**Figure 2 Legend:**
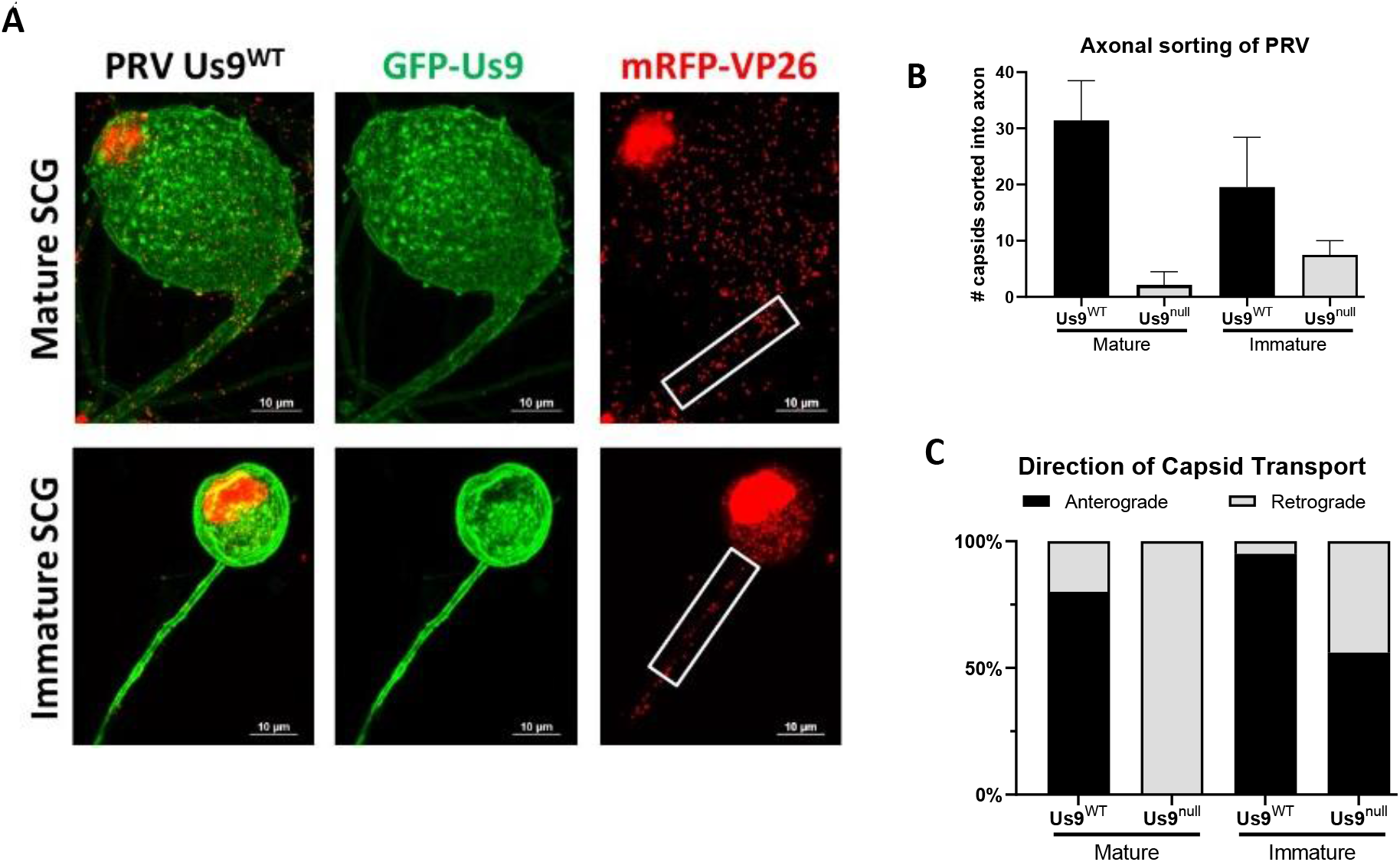
Axonal Sorting depends on Neuronal Maturation. **A**: Confocal image of SCG neuron infected with PRV expressing mRFP-VP26 (capsid) and GFP-Us9 at 10 MOI for 12 hours. The number of PRV particles, represented by mRFP-VP26 puncta, that sorted into the proximal 30um of axon (white box) are measured. **B**: Quantification of particles sorted into immature and mature SCG axons. Bars represent standard deviation. **C**: Live-microscopy quantification measuring the dynamics of particle sorting. Sorted particles were categorized as moving in the anterograde direction (away from cell-body) or retrograde direction (towards cell body).

After infection of immature SCG neurons with a dual color recombinant expressing GFP-Us9 ^WT^, the number of particles sorted into axons was slightly lower, an average of 20 particles **(Figure 2B)**. This result indicates that immature neurons are less robust compared to mature neurons at sorting of virus particles. Infection of immature SCG neurons with dual color recombinants expressing mutant GFP-Us9^YY^ had little effect Us9-dependent axonal sorting of PRV. While Us9^YY^ infection of mature SCG neurons showed a 90% reduction in particle sorting into axons, infection of immature SCG neurons reduced sorting by only 60% **(Figure 2B)**. These results show that PRV particle sorting in the neurites of immature SCG neurons is less Us9-dependent than it is in mature SCG axons.

Next, we studied the dynamics of PRV particle entry and movement in the proximal segments of mature axons or immature SCG neurites **(Figure 2C)**. All moving particles were categorized as moving away from the cell body (the anterograde direction) or towards the cell body (the retrograde direction). In dual color virus infections of mature SCG neurons expressing wild type GFP-Us9, essentially all the particles moved in the anterograde direction, consistent with previous observations (21). In the dual color infection expressing the missense GFP-Us9^YY^ protein, while fewer particles were moving, essentially all were moving in the retrograde direction. The data suggests that the retrograde moving particles are likely particles that infected axons and were in the process of moving toward the cell body and are not progeny particles.

In immature SCG neurons infected with dual color recombinants expressing wild type GFP-Us9, particle movement was predominantly in the anterograde direction, similar to that found for particles in mature neurons. This observation suggests that immature neurons can sponsor anterograde sorting and transport of PRV particles. Infection with dual color recombinants expressing mutant GFP-Us9^YY^ in immature SCG neurons **(Figure 2C)**, suggesting that virus transport in immature neurons is not completely dependent on Us9. These results suggest that PRV axonal transport is efficient in mature neurons and is unregulated in immature neurons.

### Characterization of Proteome Changes during Neuronal Maturation

The differences in PRV axonal sorting between immature and mature SCG neurons could reflect different neuronal transport mechanisms between the two developmental stages. Understanding the differences requires the characterization of how the transport-associated proteome differs with neuronal development. To test whether the differences in PRV sorting phenotypes could be due to the differences in the proteomes of immature and mature SCG neurons, we used a quantitative mass-spectrometry approach based on tandem mass tagging (TMT) to define changes in the SCG proteome with maturation. This analysis led to the identification of 4,901 quantifiable (2 peptides per protein in both replicates) proteins **(Figure 3A and Supp. Table 1)**. As expected, the neuronal maturity markers NaV1.2 and pNfH (16–18) **(Figure 1C and 1D)** were detected with higher abundances in mature samples compared to immature samples, providing confidence to the workflow **(Figure 3B)**.

**Figure 3 Legend:**
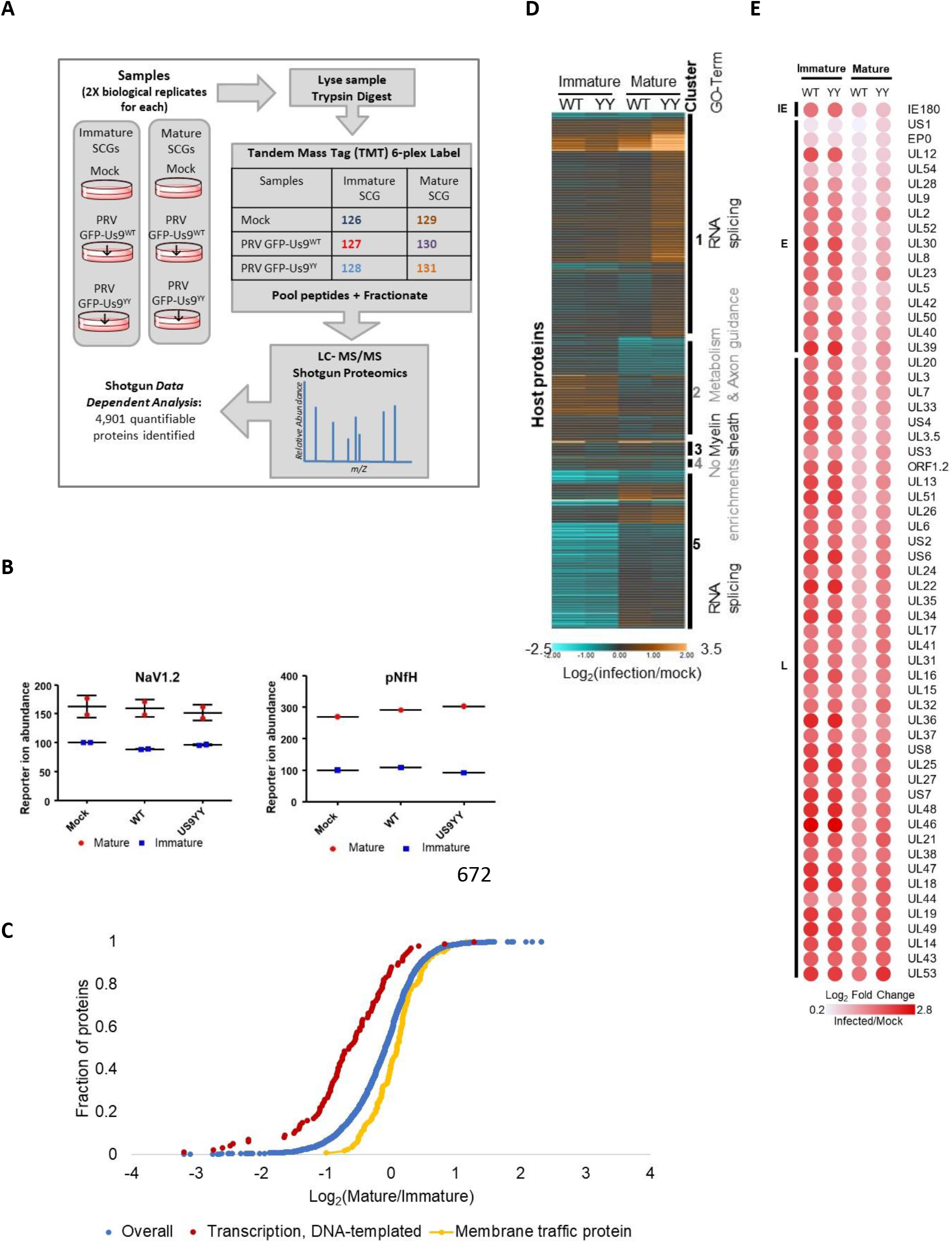
SCG Proteome Varies with Age. **A**: The workflow of the TMT mass-spectrometry experiment. **B**: Maturity markers, pNfH (top) and Nav1.2 (bottom) are detected with higher abundances in mature SCG neurons. **C**: Gene Set Enrichment Analysis of the whole proteome (blue) reveals that immature SCG neurons are enriched (FDR 2.21×10^-12^) in transcription factors (red) and mature neurons are enriched (FDR 1.38×10^-7^) in membrane-trafficking associated proteins (yellow). **D:** Host proteins altered by infection. The heatmap graphs the Log2 fold-change of host protein abundance values. Values are normalized to mock signal of the same age. All identified host proteins, that were found to be significantly differential (adjusted P-value ≤0.05) in a background-based ANOVA analysis in at least one comparison, were clustered with k means = 7. Clusters are labeled with corresponding GO-term enrichment. **E:** Viral proteins are more abundant in immature SCG neurons. The heatmap represents TMT reporter ion log2 fold-change values for PRV proteins. PRV proteins are temporally organized as IE (immediate early), E (early) and L (late) expressing.

Gene Set Enrichment Analysis for GO-term categories revealed that transcription-factor protein abundances are enriched in immature SCGs while membrane-trafficking proteins abundances are enriched in mature SCGs (**Figure 3C)**. SCG neurons, like many other neuronal subtypes, rely on membrane-trafficking proteins to allow the transport of vesicles along the axons and promote neuronal communication. The abundance of trafficking-proteins in mature neurons may explain the observed regulation of particle transport in these cells.

Additionally, we utilized the dataset to explore other functional categories affected by infection and/or neuronal age **(Figure 3D and Supp. Fig. 3 E-F)**. For example, the abundances of RNA-splicing associated proteins were affected by neuronal age but not the state of PRV Us9^WT^ or Us9^YY^ infection. Infection also increased the abundance of metabolism-associated proteins in immature neurons while the opposite was observed in mature neurons. In contrast, myelin-sheath associated protein abundances increased with Us9^WT^ infection and decreased with Us9^YY^ infection **(Figure 3D)**. Together, these observations reveal global changes to the neuronal proteome caused by maturation and/or infection and present many hypotheses underling neuronal spread of alpha-herpesvirus infections.

Within the host proteins, a small subset seems to be enriched in all infected samples, suggesting a general virus-induced neuronal response **(Supp. Fig. 3F)**. Several proteins within that group, including Annexins (Anx-1, -2, -8, -13), Got2, Myof are implicated in phospholipid-binding and cell-signaling pathways (22). Interestingly, these pathways play a role in lipid-raft related mechanisms (23), further elucidating the role of lipid-rafts in Us9-mediated anterograde spread.

To further understand the global impact of neuronal age on the progression of virus infection, virus protein abundances were monitored and represented as a heatmap **(Figure 3F)**. This revealed that both immature and mature neurons can support infection, as viral proteins were readily detected in both neuronal ages. Immature neurons however had higher abundances of virus proteins compared to the slightly muted levels in mature neurons, suggesting that mature neurons better regulate virus expression.

### Us9 forms distinct interactions in immature and mature neurons

To identify the specific proteins facilitating Us9-mediated PRV axonal sorting, we used a co-immunoaffinity purification (IP) and mass spectrometry approach **(Figure 4A)**. Mature and immature SCGs were infected with PRV-341 (GFP-Us9^WT^, mRFP-Vp26) or PRV-442 (GFP-Us9^YY^, mRFP-Vp26) for 12 h and collected in detergent-resistant-membrane preserving buffer to maintain the functional lipid-raft environment of Us9 (9, 11). GFP-based immunoaffinity enrichment coupled to data-dependent mass spectrometry was then performed to identify interacting proteins. The specificity of the interactions was assessed using the Significance Analysis of Interactome (SAINT) algorithm (24, 25), and a SAINT threshold of 0.85 (**Supp. Fig. 4A**) was employed for filtering Us9-associated proteins **(Supp. Table 2)**.

**Figure 4 Legend:**
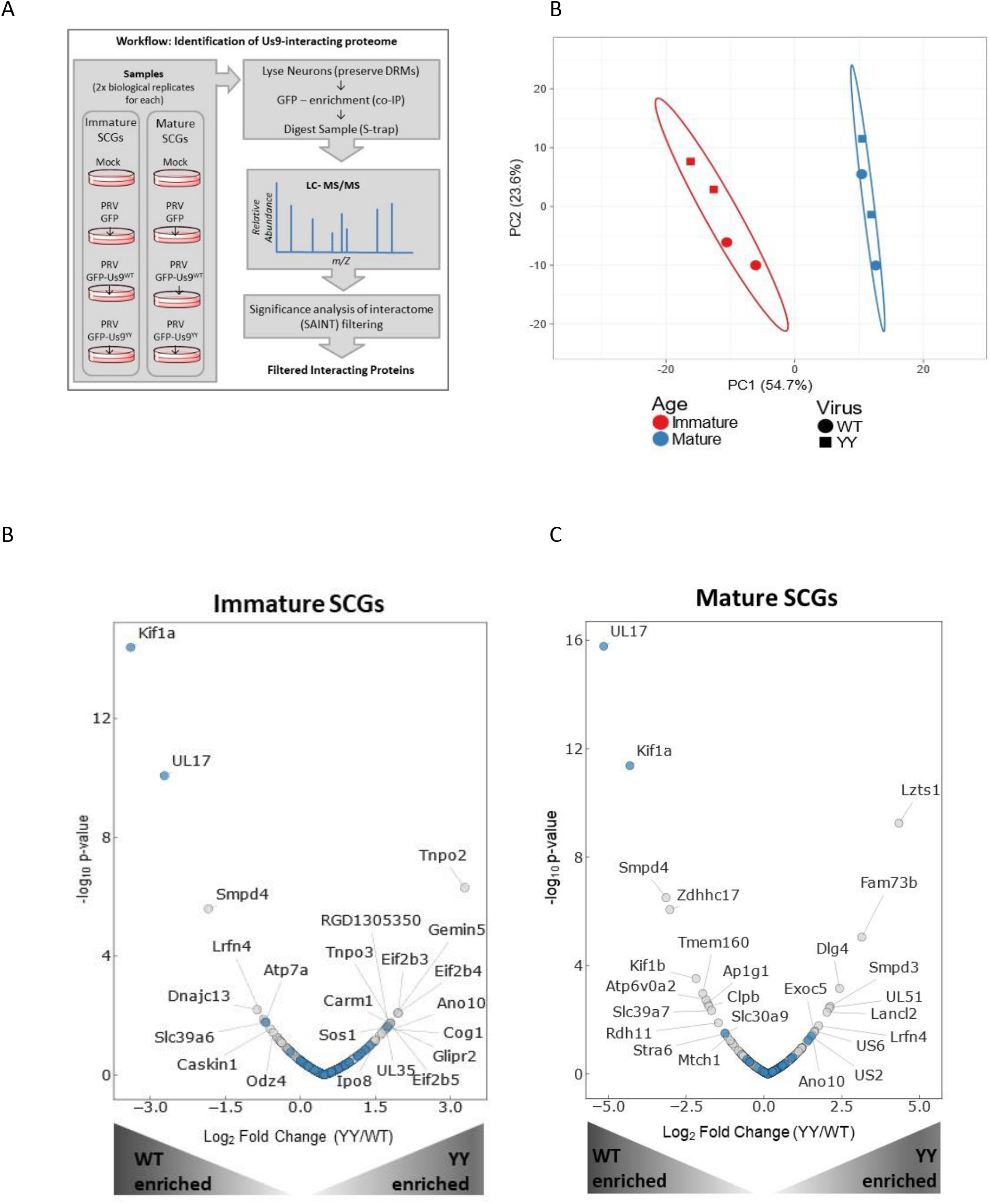
Identification of Us9-interacting neuronal proteome. **A**: Workflow describing the experimental setup. The 8 samples include immature and mature SCG neurons that are mock/uninfected or infected for 12 h with PRV 151 (GFP control), PRV 341 (GFP-Us9) or PRV 442 (GFP-Us9YY). Samples were lysed in detergent-resistant-membrane (DRM) preserving lysis buffer, followed by co-IP with GFP-conjugated magnetic beads and LC-MS/MS analysis. The resulting dataset was specificity filtered using the SAINT algorithm to identify high confidence interacting proteins. **B**: Principal Component Analysis (PCA) of the specificity-filtered data revealed clustering driven by neuronal developmental age rather than the virus state of infection. The immature neurons (blue) clustered together and the mature neurons (red) clustered together. **C and D**: Volcano-Plot representation of the immature **(C)** and mature **(D)** interactome that is associated with Us9^WT^ (left-half of plots) or Us9^YY^ (right-half of plots). Grey dots represent novel interactions and blue dots represent proteins previously reported to interact with Us9. Proteins labeled with gene names are significantly (p-value ≤0.05) differential in relative association between US9^WT^ and US9^YY^.

The identified Us9 interactions were assessed by overrepresentation analysis using GO Biological Process. The resulting treemap (**Supp. Fig. 4B**) highlighted enrichments in proteins involved in membrane transport and SNARE activities, consistent with the known role of Us9. Remarkably, a principal component analysis of the interactome revealed that the state of infection -whether mock, PRV GFP-Us9^WT^ or PRV GFP-Us9^YY^ – did not impact the Us9 interactome as drastically as neuronal age **(Figure 4B)**. This finding was supported by our observation of differential specificity for some of the interacting proteins across neuronal age (**Supp. Fig. 4C**). These results are consistent with our hypothesis that neuronal age is a predominant factor in the regulation of Us9-meadiated anterograde spread of infection.

We next examined the abundance profiles of Us9-interacting proteins to identify those that are different in Us9^WT^ versus Us9^YY^ infections **(Figure 4D)**. A volcano plot representation of the interactome data revealed that the most abundant neuronal protein associated with Us9^WT^ is Kif1a, an anterograde-directed microtubule motor that has been shown to interact with Us9 to facilitate PRV spread (9). This prominent interaction with Kif1a served as validation of our interaction study, adding confidence to the identified interactions that were not previously reported. The strong association of Us9^WT^ with the PRV protein UL17 is an intriguing finding **(Figure 4C, D).** Our study also uncovered additional Us9 interactions with cellular proteins in mature neurons. Interaction with Zdhhc17/Hip-14, a palmitoyltransferase, is interesting because of its role in neuronal transport mechanisms **(Figure 4D)**. Zdhhch17 palmytolates Snap25 (26), a SNARE protein that was previously identified in the Us9-interactome (9).

We suggest that the association of Us9^WT^ with vesicle-transport associated proteins (such as Kif1a, SMPD4, Zdhhc17, Kif1b, Atp6Voa2, Ap1g1, Slc39a&) in mature neurons indicates the presence of mechanisms that promote anterograde spread. The Us9^YY^ associated proteins may represent the lipid-raft milieu in which Us9^YY^ is present but fails to interact with machinery that facilitate transport.

### SMPD4 regulation of virus spread

We identified SMPD4/n-sMase3, a sphingomylinase found in lipid-rafts that is known to function in vesicle membrane processes, as the most abundant neuronal protein associated with Us9^WT^. To assess the role of SMPD4 in PRV spread, an siRNA knockdown followed by a tri-chamber anterograde spread assay was performed **(Figure 5A)**. The chamber allows for the physical separation of SCG neuronal cell bodies in the soma-S-compartment, and axonal termini in the neurite-N-compartment. Upon infection in the S-compartment, virus particles that undergo successful axonal sorting and anterograde spread can enter the N-compartment where they are amplified by the PK15 cells. Thus, anterograde spread can be assayed by the measurement of fluorescent virus particles and titer in the N-compartment.

**Figure 5 Legend:**
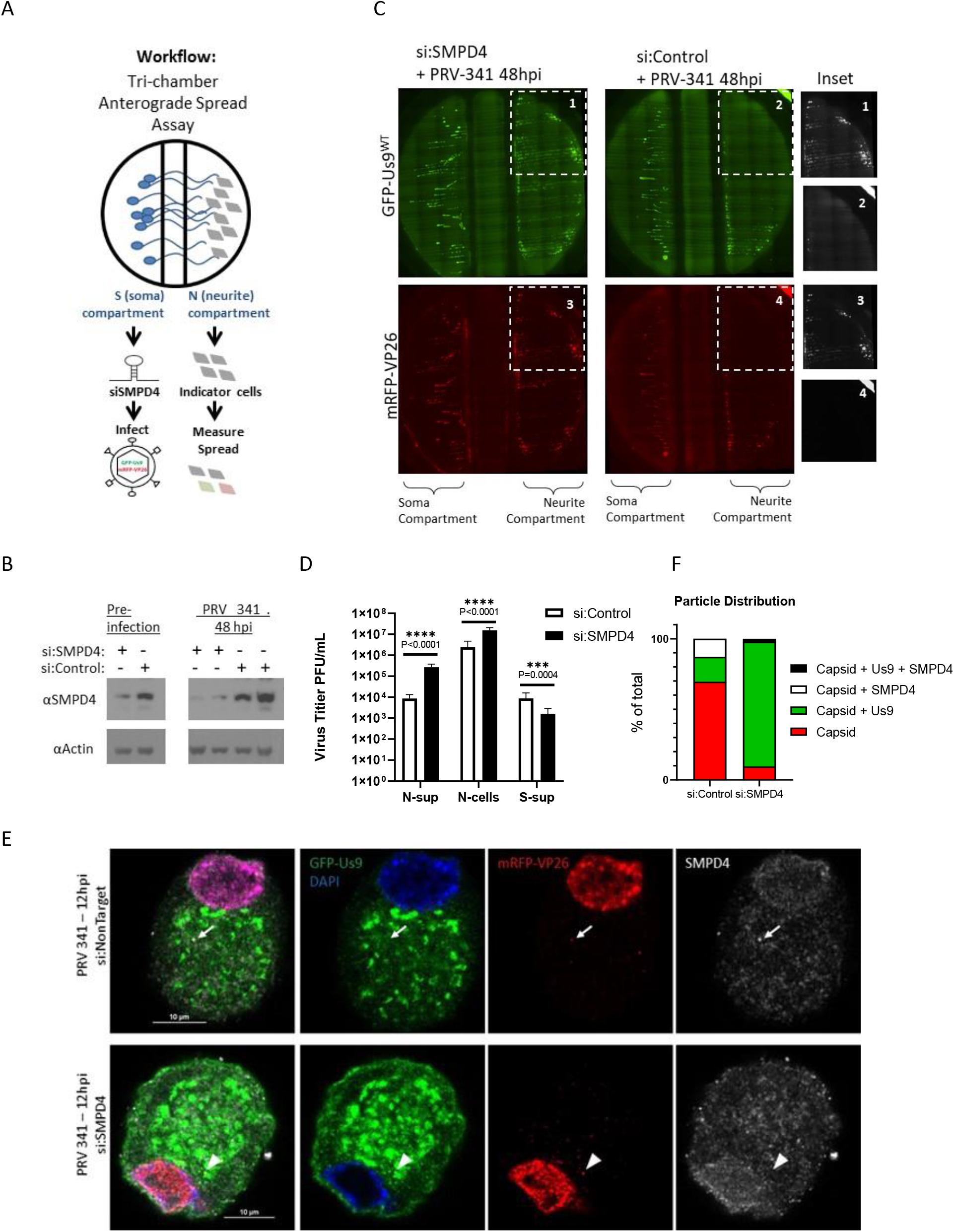
SMPD4 knockdown facilitates PRV Spread. **A**: Tri-chamber Anterograde Spread Assay workflow– Dissociated SCG neurons are seeded in the soma-S-compartment (left), growing axons penetrate through the middle-M-compartment into the neurite-N-compartment (right). siRNA are administered in the S-compartment for 3 days, followed by infection in the S-compartment. The spread of virus particles into the N-compartment can be detected by fluorescent expression of GFP-Us9 or mRFP-VP26 in the N-compartment. **B:** SMPD4 siRNA knockdown. Dissociated SCG neurons were transfected with 50uM of siRNA against SMPD4 (+) or Non-Target controls (−). At 3 days post siRNA transfection (labeled pre-infection), samples were collected and assayed on SDS-PAGE western blot to confirm protein knockdown. After the anterograde sorting assay, Soma form the S-compartment were collected again to measure knockdown for the duration of the assay. Each lane represents a different chamber. **C**: Robust spread detected after SMPD4 knockdown. At 48 hpi, the N-compartment of chamber treated with siRNA-SMPD4 (left) displayed greater GFP-Us9 (top) and mRFP-VP26 (bottom) signal, in comparison to the si:NonTarget negative-control (right chamber). **D:** Virus titers after anterograde sorting assay. N-sup represents virus particles that have sorted into the N-compartment and released into the supernatant. N-cells represents particles sorted into the N-compartment but confined inside the axons or PK-15 cells. S-Sup represent particles released into the supernatant of the S-compartment. Titer was measured by counting plaques on a monolayer of PK-15 cells. Statistics were performed using 2way-ANOVA test. **E:** SMPD4 localization after PRV infection. Confocal microscopy of siRNA transduced SCG cell body infected with PRV-341 expressing GFP-Us9 and mRFP-VP26 capsids. After 12 hpi, cells were fixed for immunofluorescence staining of SMPD4. White arrow indicates foci of mRFP-VP26 and SMPD4 colocalization. Arrowhead indicates foci of mRFP-VP26 and GFP-Us9 colocalization. **F:** Quantification of mRFP-VP26 capsid distribution. All cytoplasmic mRFP-VP26 capsid foci were quantified for colocalization with GFP-Us9 and/or SMPD4 foci.

To understand if SMPD4 was necessary for Us9-mediated anterograde sorting, the endogenous protein was reduced with siRNA **(Figure 5B)**. Dissociated SCG cell-bodies in the soma-compartment were transduced with siRNA targeting SMPD4 (si:SMPD4) or non-targeting siRNA (si:control). At 3 dpt (days post transfection), pre-infection cell lysates were collected to confirm protein knockdown, followed by PRV-341 infection at MOI-10. At 48 hpi the cell bodies in the S-compartment were collected to assay continued knockdown of SMPD4 expression at the end of anterograde sorting assay. Western blot confirms SMPD4 knockdown for the duration of the anterograde sorting assay **(Figure 5B)**.

Upon siRNA knockdown of SMPD4 in the soma-compartment, an increase in the expression of GFP-Us9 and mRFP-VP26 was detected in the neurite-compartment, suggesting an increase in anterograde spread upon SMPD4 knockdown **(Figure 5C)**. Consistently, an increase in PRV titer was also detected in the N-compartment upon siRNA knockdown of SMPD4 **(Figure 5D)**. Surprisingly, the soma-compartment titer was reduced upon SMPD4 knockdown, suggesting that virus particles are efficiently being sorted down the axon rather than being released from the cell-body **(Figure 5D)**. These results indicate that SMPD4 may be involved in counteracting PRV anterograde spread, an unexpected antiviral function.

Immunofluorescence and confocal microscopy of infected SCG neurons co-stained with SMPD4 antibody revealed several SMPD4 foci localized throughout the SCG soma **(Figure 5E and Supp. Fig. 5C)**. In mock cells, SMPD4 localization displays nuclear, nuclear-envelope, and cytoplasmic localization. While siRNA knockdown does not complexly eliminate antibody staining, both the nuclear-envelope and cytoplasmic foci are largely reduced **(Supp. Fig. 5C)**. Upon infection, some mRFP-VP26 capsids colocalized with SMPD4 foci but not GFP-Us9 (arrow). After siRNA knockdown, mRFP-VP26 capsids lost SMPD4 association but colocalized with GFP-Us9 (arrowhead) **(Figure 5E)**. Quantification of capsid association revealed that while 18% of capsids co-localize with GFP-Us9 in wildtype infections, knockdown of SMPD4 increased capsid-Us9 association to 88% **(Figure 5F)**. This further supports a role for SMPD4 in counteracting Us9-capsid association.

It has been proposed that SMPD4 expression decreases with neuronal age and that it may sensitize cells to DNA damage-induced apoptosis (27). We also observed that SMPD4 abundance decreases with neuronal maturation **(Supp. Fig. 5A)**. Additionally, we observed that increased SMPD4 expression correlates with increased LC3 expression, a marker of cellular stress and autophagy **(Supp. Fig. 5B)**. Interestingly, infection with Us9-deleted virus did not express the same levels of SMPD4 or LC3, which further supports a function with Us9 and neuronal stress-response. It will be interesting to explore how Us9 expression may stimulates SMPD4 expression, perhaps Us9 plays an unknown role in stimulating DNA damage and/or apoptosis.

## Discussion

Alpha-herpesviruses establish a life-long latent infection in peripheral neurons that can sporadically reactivate over the lifetime of the host and lead to transmission of the infection to other hosts. The spread and pathogenesis of alpha herpesvirus infections are affected by the age of the infected host, but the underlying mechanisms are unknown. This study examined how the stage of neuronal development affects the spread of PRV infection and its dynamics in cultured neurons. We identified several neuronal proteins that may facilitate spread of infection. One of these, the lipid raft protein SMPD4, had an unexpected function and may have a unique antiviral role in regulating axonal sorting and anterograde spread.

We established and characterized primary neuronal cultures of immature and mature rodent superior cervical ganglia (SCG), sympathetic neurons of the peripheral nervous system. Maturity was defined by age in culture, generation of an extensive axonal network and the appearance of neuronal proteins typical of terminally differentiated neurons. Our proteomics experiments revealed substantial differences between immature and mature SCG neurons. The higher abundance of membrane trafficking associated proteins in mature neurons and their lack in immature SCG neurons supports our hypothesis and model that PRV anterograde spread is efficient and robust in mature neurons while it is non-specific and unregulated in immature neurons **(Figure 6A)**.

**Figure 6 Legend:**
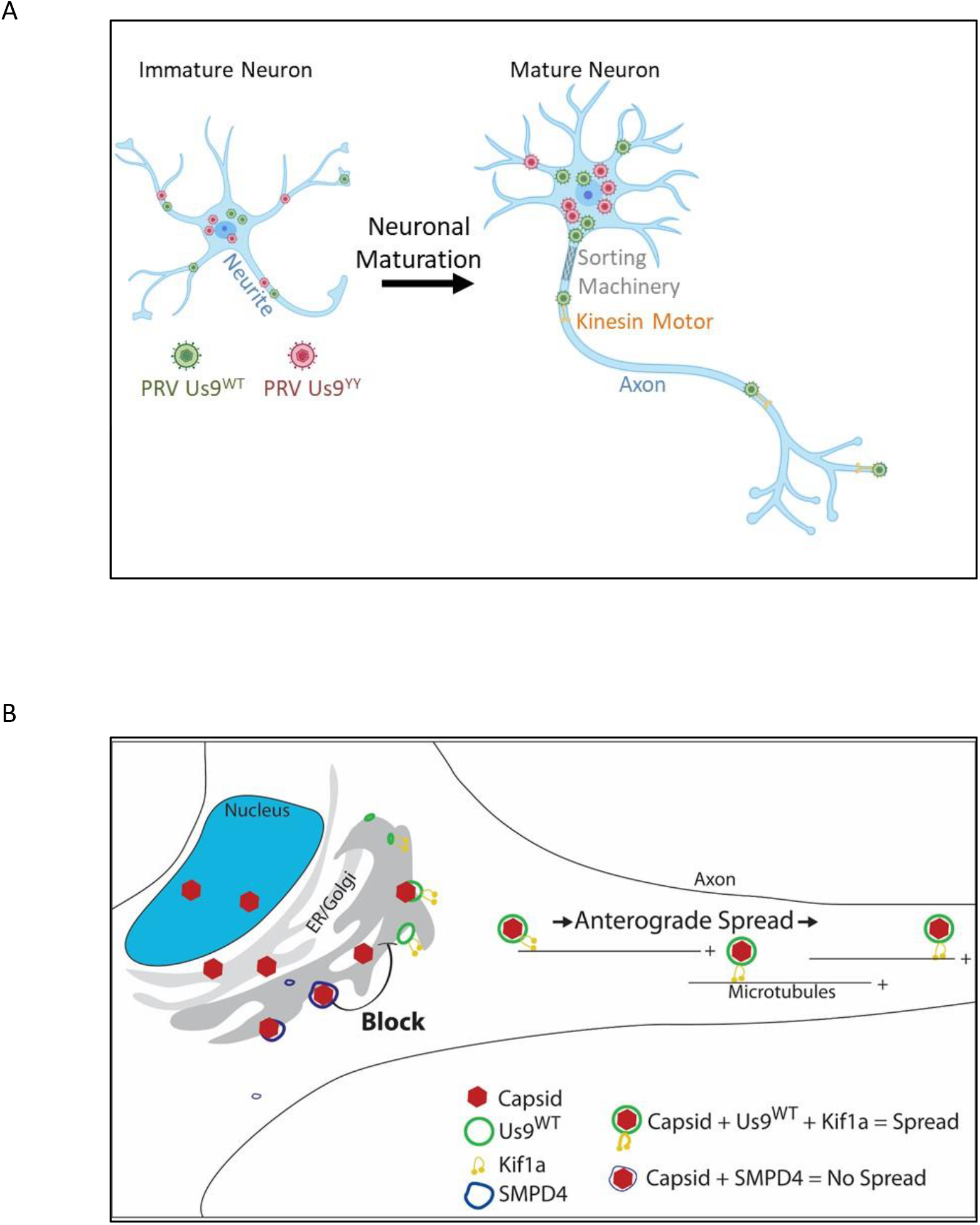
Model. **A:** Neuronal Maturation is required for efficient and robust anterograde spread. Immature neurons lack the proteome necessary to regulate spread of virus particles. PRV particles expressing Us9^WT^ or the spread deficient Us9^YY^ can sort. Neuronal maturation is accompanied with establishing an Axon and expression of proteins specialized to regulate anterograde spread. PRV particles expressing Us9^WT^ spread but not Us9^YY^. **B:** SMPD4 blocks anterograde spread. Capsids colocalizing with SMPD4 foci (blue) do not colocalize with Us9 and do not recruit transport machinery, such as the Kif1a microtubule motor, to facilitate anterograde spread along the axon.

These data established the marked differences that occur as neurons differentiate and highlight the importance of time in culture for functional studies. After sufficient time in culture (such as 20 days), mature SCG neurons are more uniform in their proteome and function, while immature cultures are a mixture of cells in different stages of growth and maturation. As a result, there is more variability in almost any measurement when immature neurons are analyzed.

Our studies focused on a unique and essential aspect of alphaherpesvirus infection of neurons, the sorting of viral particles into axons and anterograde spread of infection in the nervous system. For PRV, such sorting is dependent on the viral Us9 protein. Previous work in our lab had found that sorting of PRV structural membrane proteins into immature axons was not dependent on the Us9 protein [17]. We extended this observation to show that mature neurons sponsored robust and efficient axonal sorting and anterograde spread of Us9-medated PRV particles. Immature neurons, on the other hand, were less dependent on Us9 for sorting into neurites. These immature structures have not developed into mature axons and do not produce the protein composition that specializes in vesicle-transport and trafficking mechanisms **(Figure 6A)**.

By using Us9-GFP fusion proteins, we used immunoaffinity co-purification to identify Us9 interacting proteins. Our study identified several previously unknown Us9-interactors in mature SCG neurons and confirmed other previously identified Us9 interacting proteins. In particular, we detected the Kif1a kinesin motor as a Us9 interacting protein in mature SCG neurons, which supports the previous findings of Kramer *et al*, who found this interaction in PC12-cell cells. We also identified Kif1b, a paralog of Kif1a as a potential motor facilitating PRV anterograde spread (28). We found an interaction between Us9 and Zdhhc17, a palmitoyltransferase of the SNARE protein Snap25, implicated in the regulation of neuronal vesicle-trafficking and associated with neurological diseases such as Huntington’s (26, 29). Previous studies proposed an interaction of Us9 with Snap25 (9), and preliminary experiments suggest that Us9 and capsids co-transport with Snap25 in vesicles.

We were surprised to see a significant interaction of Us9 with viral protein UL17. To date, all known functions of UL17 involve viral DNA encapsidation, capsid formation and maturation in the nucleus (30, 31). In contrast, all known Us9 functions reside outside of the nucleus. This interaction suggests a potential new role for UL17 outside the nucleus where it may interact with Us9. Preliminary experiments investigating the significance of this interaction are underway.

We made particular use of recombinant viruses that expressed a Us9 missense protein, which is a functional null mutant. It is found in lipid rafts like the wild type Us9 protein, but it does not interact with Kif1a and does not sponsor sorting of particles into axons. Using the GFP-Us9 missense protein, we could differentiate proteins within lipid-rafts milieu that may facilitate Us9-dependent anterograde spread from those that simply are present in the raft milieu.

While we anticipated that the Us9^WT^ interacting proteins would be those that facilitated PRV anterograde spread, we made an unexpected finding, a negative regulator of Us9 function. SMPD4 was one of the most abundant interactions with GFP-Us9. It is a sphingomyelinase found in lipid rafts. SMPD4 catalyzes hydrolysis of sphingomyelin into phosphorylcholine and ceramide and plays a crucial role in the lipid-rafts by regulation membrane composition (32). Lipid-rafts have an important role in neuronal function and in the pathogenesis of many viruses, including herpesviruses and PRV (11). The association of PRV Us9 with SMPD4 is interesting, as both are constituents of lipid-rafts and are involved in membrane-trafficking processes (11, 33). Genetic defects with Smpd4 inheritance present cognitive problems, defects in brain development and microcephaly (27), thus further studies may illuminate a relationship between neurotropic infections and such disease outcomes. We found that when SMPD4 is reduced, PRV anterograde neuronal spread is increased, rather than decreased. These results support a model in which SMPD4 negatively regulates PRV axonal sorting and subsequent anterograde neuronal spread by preventing the association of capsids with Us9-associated transport vesicles **(Figure 6B)**. In wildtype infection, SMPD4 colocalizes with capsids and may reduce capsid association with Us9, which is required for anterograde spread. Upon knockdown, capsids colocalize with Us9 and undergo anterograde spread. Together, these results suggest an inhibitory, anti-viral function for SMPD4 in PRV infection. A further test of the antiviral role of SMPD4 would be to know if the widely used statins, drugs that reduce cholesterol, affect the spread of PRV or other neurotropic viruses as it has been shown for Ebola (34).

## Materials and Methods

### *Ex vivo* primary cultures of SCGs

Sprague Dawley Rattus norvegicus were purchased from Charles River Laboratories (Maryland, USA) at E16.5. Embryos were harvested on E17.5 for dissection of the Superior Cervical Ganglia. The ganglia are dissociated and seeded on Poly-ornithine treated and laminin-coated tissue-culture dishes. Neurons are suspended in neurobasal media supplemented with B27 and NGF (35). At 2 DIV (days in vitro) the neurons were treated with AraC, an antimitotic drug, to eliminate the contaminating non-neuronal cells. Neurons were collected in PBS for analysis at 4 DIV as immature cells, and 20+ DIV for mature neurons. Ethics Statement: All animal work was performed as approved by the Princeton Institutional Animal Care and Use Committee (IACUC) protocol 1947. All Princeton University animal work personnel are required to adhere to local, state and federal laws governing animal research and is accredited by the Association for Assessment and Accreditation of Laboratory Animal Care (AAALAC).

### Protein Quantification with BSA

Neuronal samples were collected in PBS at appropriate DIV, spun down at 4C for 5min at 13K, the PBS supernatant was discarded, and pellet was retained. BSA Assay per manufacturer’s instructions were followed.

### Virus Infections

Pseudorabies Virus stocks are grown and tittered on PK15 cells. SCG neurons are infected at a high MOI of 10, neurons are incubated for 1 hour after which virus inoculum is replaced with neuronal media. Virus strains used in this study include: PRV-341 (GFP-Us9^WT^, VP26-mRFP), PRV-442 (GFP-Us9^YY^, VP26-mRFP), PRV-151 (diffusible GFP) (20, 36).

### Axonal Sorting Assay

At 12 hpi (hours post infection), infected cultures were fixed with 4% para-formaldehyde followed by 2X-PBS washes. Brightfield microscopy was performed with Nikon Eclipse-Ti instrument. NIS-Elements AR V4.3 was used to capture and analyze images. The number of sorted particles was acquired by counting the number of mRFP-VP26 foci in the proximal 30um of axons.

### Immunofluorescence microscopy

Neurons are fixed in 4% Paraformaldehyde-PBS solution for 10 minutes at room temperature, followed by 2X washes in PBS, then permeabilization in 0.5% Trypsin-PBS for 20 min at room temperature (RT) followed by 2x PBS washes. Cells were blocked with 3% BSA-PBS for 1 h followed incubation in primary antibody for 2 h at RT, then 2X PBS, followed by a secondary antibody for 1h RT and then 2x PBS washes. Confocal images were acquired on Nikon Ai1R with NIS-Elements Ar-v4.5.

### Quantification using Tandem Mass Tagging (TMT) Mass Spectrometry

#### Cell lysis, protein digestion and TMT labeling

Infected cells were lysed in 100 mM Tris-HCl, pH 8.0, 2% SDS, 1 mM EDTA preheated to 70 °C. Cell pellets were subjected to three successive rounds of heating at 95 °C and sonication. Protein concentration was assessed by BCA assay (Pierce) and 100μg of protein from each sample was reduced with 25 mM tris(2-carboxyethyl)phosphine (Pierce) and alkylated with 50 mM chloroacetamide at 70 °C for 20 min. The protein was then precipitated via methanol-chloroform cleanup (37). Samples were digested overnight at 37 °C in 50 mM HEPES pH 8.3 at a 1:50 trypsin:protein ratio (Pierce). Peptides from both biological replicates were labeled with a 6-plex TMT kit as previously described (38) and pooled in equal peptide amounts, resulting in two individual 6-plex experiments. Pooled peptides were fractionated by basic pH reverse phase C18 StageTips. After binding peptides to the C18 material, a wash with 5% ACN was performed, followed by step-wise elution using a gradient of 8% - 46% ACN in steps of 2%, resulting in 20 fractions. The 20 fractions were concatenated into 10 final fractions by combining fractions 1 and 11, 2 and 12, etc. Fractionated peptides were dried in vacuo and resuspended in 5 μL of 1% FA, 2% ACN in water.

#### LC-MS/MS and bioinformatic analysis

Peptides (2 μL) were analyzed via LC-MS/MS using a Dionex Ultimate 3000 UPLC coupled online to a Nanospray Flex ion source and a Q Exactive HF. Reverse-phase chromatography was performed at 50 °C over a 25 cm IntegraFrit column (IF360–75-50-N-5, New Objective, Woburn, MA) packed in-house with 1.9 μm ReproSil-Pur C18-AQ (Dr. Maisch, GmbH) with mobile phase A: 0.1% formic acid in water and mobile phase B: 0.1% formic acid in 97% acetonitrile. A 120 min gradient consisting of 4% B to 12% B over 60 minutes, followed by 12% B to 25% B over 60 min was used to separate the peptides. Following ionization at 2.1kV, an MS1 survey scan was performed from 350 to 1800 m/z at 120,000 resolution with an automatic gain control (AGC) setting of 3e6 and a maximum injection time (MIT) of 30 ms recorded in profile. The top 20 precursors were then selected for fragmentation and MS2 scans were acquired at a resolution of 30,000 with an AGC setting of 1e5, a MIT of 50 ms, an isolation window of 0.8 m/z, a fixed first mass of 100 m/z, normalized collision energy of 34, intensity threshold of 1e5, peptide match set to preferred, and a dynamic exclusion of 45 s recorded in profile.

MS/MS data were analyzed by Proteome Discoverer (Thermo Fisher Scientific, v2.2.0.388). A fully tryptic search against a combined mouse, rat, and pseudorabies virus Uniprot database appended with common contaminant sequences (downloaded 3/2017 – 80,004 sequences) requiring 4 ppm mass accuracy on the precursor ions and 0.02 Da accuracy on the fragment ions was performed. Static carbamidomethyl modifications to cysteine, static TMT additions to peptide N-termini and lysine residues, dynamic oxidation of methionine, dynamic deamidation of asparagine, dynamic methionine loss and acetylation of protein n-termini, and dynamic phosphorylation of serine, threonine, and tyrosine were allowed as modifications in the search. Matched spectra were scored by Percolator and reporter ion signal-to-noise values were extracted. Following parsimonious protein assembly at a 1% FDR for proteins and peptides, reporter ion quantitation was performed for unique and razor peptides with an average signal/noise (S/N) ratio of at least 8 and a precursor co-isolation threshold of less than 30% which did not contain a variable modification and normalized to the total detected signal in each TMT channel. Protein abundances were calculated as the sum of all reporter ion values in a particular channel for each protein. Imputation of missing values was performed by low abundance resampling. The data were scaled based on the Immature Mock infection samples. Statistically differential proteins were assessed via the background based ANOVA implemented in Proteome Discoverer. The resulting data was exported to Excel for further analysis. Individual protein graphs were made using Graphpad Prism, v5.04. Gene Set Enrichment Analyses were performed using Pantherdb.org.

### Immunoaffinity purification-Mass Spectrometry (IP-MS) Method

#### Cell lysis, IP, and protein digestion

Cells were lysed using a previously optimized buffer for Us9 (20 mM HEPES-KOH [pH 7.4], 110 mM potassium phosphate, 2 mM ZnCl2, 0.1% Tween-20, 1% Triton X-100, 150 mM NaCl, and protease inhibitor cocktail [Sigma-Aldrich, Saint Louis, MO] at 1:100) (9). Following addition of lysis buffer, cell pellets were homogenized by Polytron (Kinematica) for 20s at 20,000 rpm. Lysates were then pelleted at 10,000 × g for 10 min at 4 °C. Clarified lysates were then added to GFP-Trap MA beads (gtma-100, Chromotek, Hauppauge, NY). For each IP, 20 μL of bead slurry was washed 3 × 500 μL in wash buffer (lysis buffer without inhibitors and nuclease). Soluble lysates were added to the beads and incubated for 60 min at 4 °C with end-over-end rotation. Following the incubation, the beads with bound proteins were collected via a magnetic rack and then suspended in wash buffer and transferred to a new tube. The beads were then washed 3 × 500 μL in wash buffer with magnetic collection in between each wash and then resuspended in 500 μL H2O and transferred to another tube. The beads were washed a final time with H2O and then eluted in 50 μL of 106 mM Tris HCl, 141 mM Tris Base, 2% SDS, 0.5 mM EDTA. Elutions were then reduced to 20 μL volume via vacuum centrifugation, and reduced and alkylated with 25 mM TCEP (77720, Thermo Fisher Scientific) and 50 mM chloroacetamide respectively at 70 °C for 20 min. The elutions were then digested via S-Trap (Protifi) according to the manufacturer’s instructions using the high recovery protocol with a one-hour digest.

#### LC-MS/MS and bioinformatic analysis

Peptide samples were analyzed on an Ultimate 3000 nanoRSLC coupled online with an ESI-LTQ-Orbitrap Velos ETD mass spectrometer (Thermo Electron, San Jose, CA). Reverse-phase chromatography was performed over a 20 cm IntegraFrit column (IF360–75-50-N-5, New Objective, Woburn, MA) packed in-house with 1.9 μm ReproSil-Pur C18-AQ (Dr. Maisch, GmbH) with mobile phase A: 0.1% formic acid in water and mobile phase B: 0.1% formic acid in 97% acetonitrile. Peptides were separated over a 150 min gradient (5% B to 30% B) with 250 nl/min flow rate and analyzed by MS1 survey scans followed by data-dependent collision-induced dissociation (CID) MS/MS fragmentation of top 15 most abundant ions. The following parameters were used: FT preview scan disabled, waveform injection and dynamic exclusion enabled, automatic gain control target value of 1 × 10^6^ for MS and 1 × 10^4^ for ion trap MS/MS scans, max ion injection time of 300 ms for MS and 125 ms for MS/MS scans. For MS scans: m/z range of 350– 1700 and resolution of 120,000; for MS/MS scans: minimum signal of 1,000, isolation width of 2.0, normalized collision energy of 30% and activation time of 10 ms.

MS/MS spectra were searched against a combined mouse, rat, and pseudorabies virus Uniprot database appended with common contaminant sequences (downloaded 3/2017 – 80,004 sequences) using Proteome Discoverer 2.2.0.388. The Spectrum Files RC node and Minora Feature Detector nodes were used to perform offline mass recalibration and label-free MS1 quantitation respectively.

The data were searched using Sequest HT with settings for a fully tryptic search with a maximum of two missed cleavages, precursor mass tolerance of 5 ppm, fragment mass tolerance of 0.3 Da, static carbamidomethylation of cysteine, dynamic oxidation of methionine, dynamic deamidation of asparagine, and dynamic loss of methione plus acetylation of the protein N terminus. Matched spectra were scored by Percolator. Label-free MS1 quantitation was performed using the max peak intensity for each peptide. For protein inference, two unique peptide sequences were required, and parsimonious assembly was performed. Only unique and razor peptides were used for MS1 quantitation. Data were exported to excel for further analysis. Proteins with at least 8 spectra identified across the entire dataset were considered for further analysis.

Total spectral count data was analyzed by SAINT (24) using the REPRINT (25) interface. SAINT was run with LowMode off, MinFold on, and Normalize on and the average SAINT score in each condition was used for specificity assessment. Based on the distribution of the SAINT scores and the identification of previously-known Us9 interactions, a SAINT threshold of ≥ 0.85 was selected. Proteins passing specificity thresholds were further analyzed using MS1 abundance-based quantitation. Principal component analysis (PCA) was conducted using Clustvis (39). Individual protein graphs were made using Graphpad Prism, v5.04. Gene Set Enrichment Analyses were performed using Pantherdb.org. Volcano plots were generated using Instant Clue (40). The mass spectrometry proteomics data reported in this paper have been deposited at the ProteomeXchange Consortium via the PRIDE partner repository (41). The PRIDE accession number is PXD 017822.

### Tri-chamber Anterograde Spread Assay

Dissociated SCG neurons are seeded in the S-compartment of campenot tri-chambers and cultured for a minimum of two weeks to allow axons to penetrate the N-compartment. Further details of this method are described in Curanovic et al (20, 35). After 14 days, PK15 indicator cells are seeded on top of axons in the N-compartment and SCG soma in the S-compartment are infected with PRV-341 virus (strain expressing GFP-Us9 and mRFP-VP26) at 10 MOI. The chamber is imaged by fluorescent microscopy every 12 hours, up to 48 hours post infection.

### siRNA knockdown

After 14 DIV in the Tri-chamber, neuronal soma in the S-compartment are transduced with siRNA (Dharmacon). 100nM of siRNA for SMPD4 or NonTarget negative-control are transfected according to manufacturer protocol by magnetofection (OZbiosciences). Samples were collected in 2X-Laemmli buffer to confirm knocked at either 3 dpt (days post transfection) or after completion of the Tri-chamber anterograde sorting assay.

## Acknowledgements

This research was supported by funding from the NIH NIGMS (GM114141) to I.M.C., and the NIH NIGMS (T32GM007388) to N.S.T.

---

**Supplemental 3 Legend:**
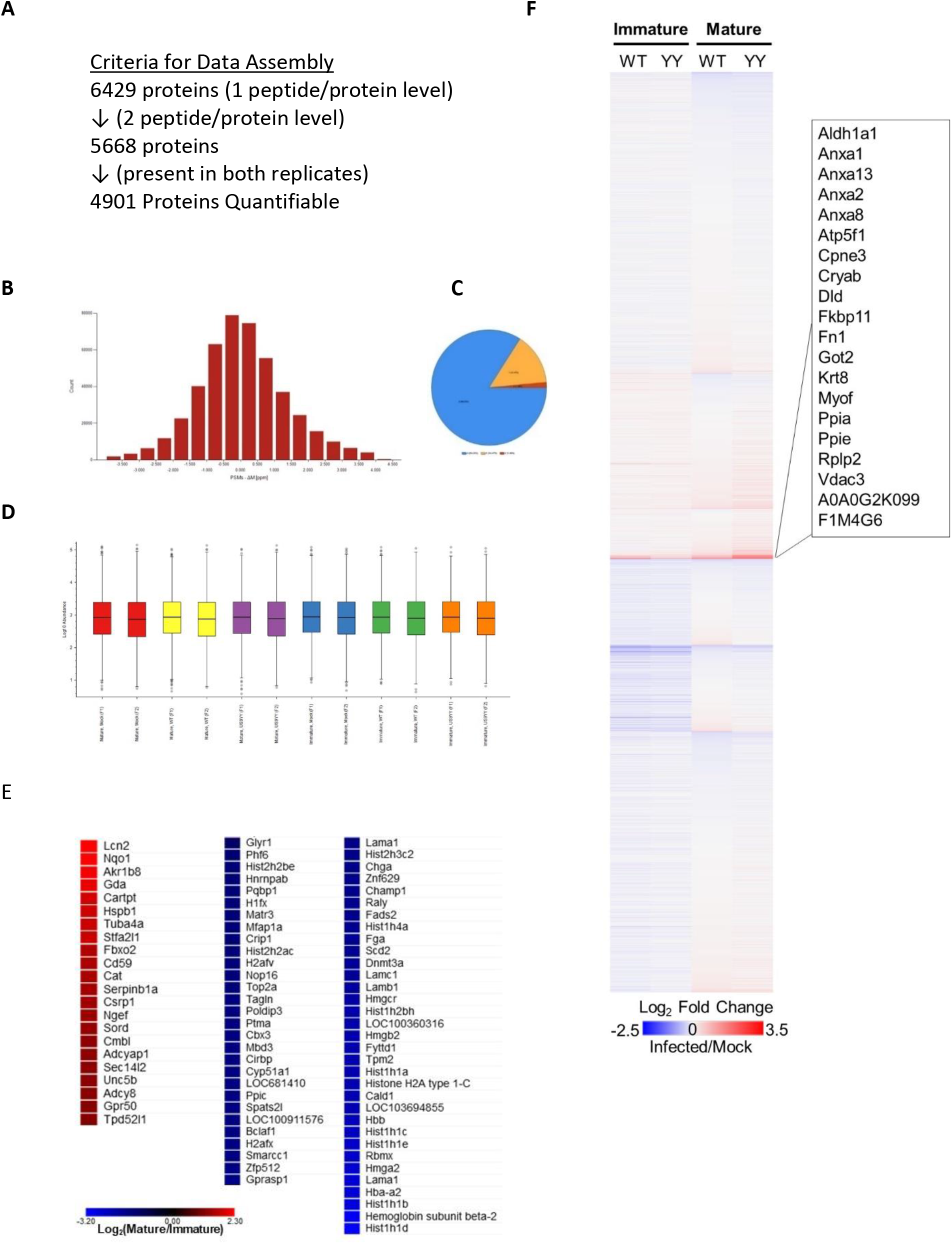
Tandem Mass Tag mass spectrometry Analysis. **A**: Criteria used to filter the data set. **B**: Mass Accuracy: High mass accuracy centered about 0ppm is reliable. **C:** Missed Cleavages - Relatively low missed cleavage rate demonstrating thorough trypsin digest of peptides. **D**: Sample Abundances - Equal abundances across all sample suggests comparable mixing between samples. **E:** Differences in mock mature v immature proteome. These proteins were significantly differential by background-based ANOVA analysis and are ordered by abundance levels (higher in Mature on upper left and higher in immature in bottom right). Chromatin organizing proteins are overrepresented here (adjusted P-value <7.68E-10). **F:** TMT reporter ion values for host proteins. Many proteins express similar levels. A small subset appears enriched in all infection conditions; the individual protein names are listed in the inset.

**Supplemental 4 Legend:**
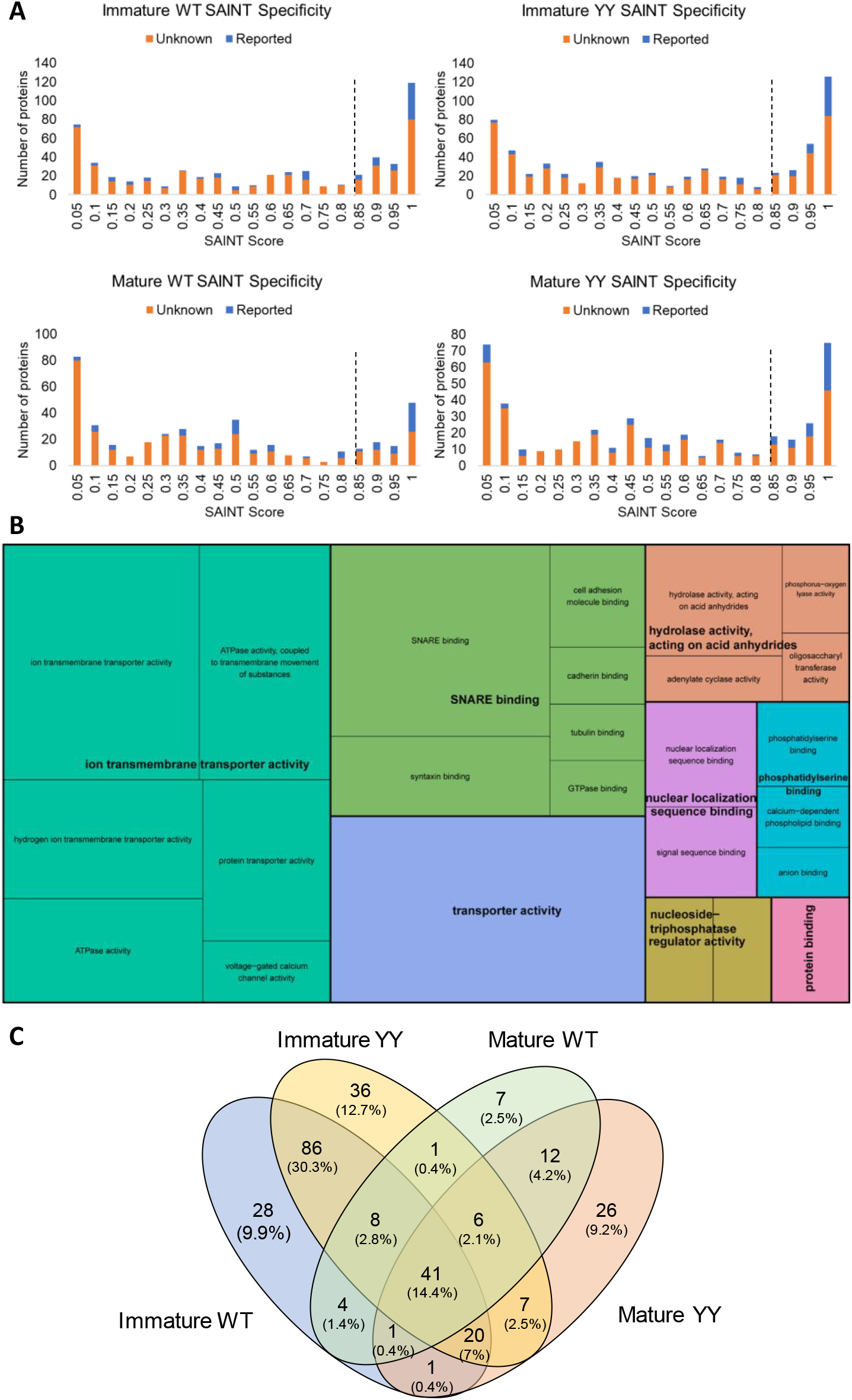
Analysis of US9 IP-MS. **A**: SAINT cutoff (dashed line) of ≥0.85 chosen for Us9 baits based on histogram of SAINT scores for novel (orange) and previously reported (blue) interactions. **B**: Treemap of Enrichment analysis by GO (gene ontology) MF (molecular function) terms. Functionally related categories are grouped by color and boxes are sized by adjusted p-value. Enrichment analysis of all specificity-filtered Us9 interacting proteins shows enrichment in transporter and SNARE binding activity. **C**: Venn diagram of specificity-filtered proteins in Us9^WT^ and Us9^YY^ IPs highlights that some proteins are found as interactors in both neuronal developmental stages and some are unique to one stage.

**Supplemental 5 Legend:**
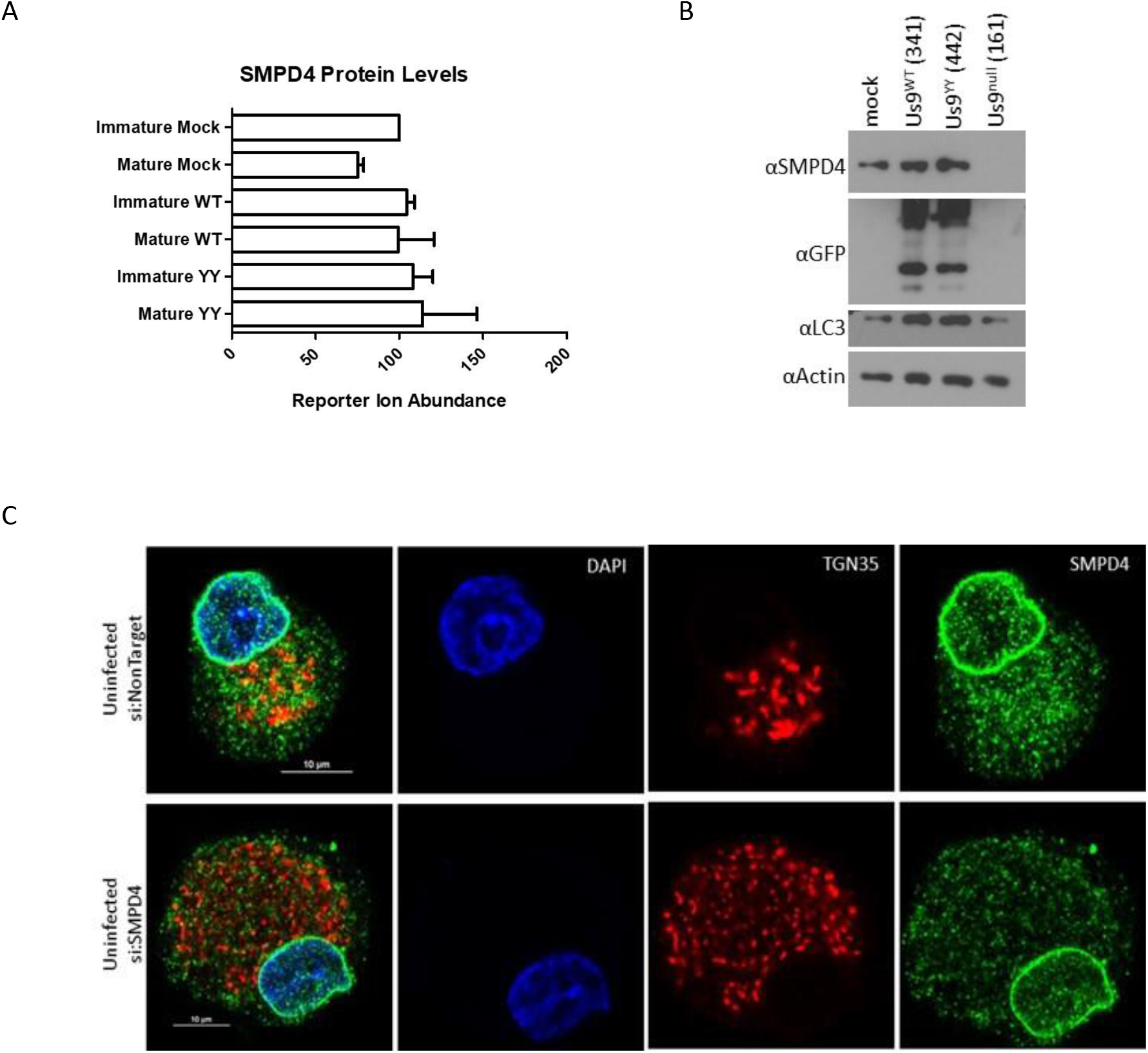
SMPD4 Characterization. **A:** SMPD4 protein abundance values from mass-spectrometry. In uninfected/mock dissociated SCGs, SMPD4 is slightly more abundant in mature neurons compared to immature. Infection increases expression levels compared to mock, but SMPD4 levels do not change between Us9^WT^ or Us9^YY^ infection conditions. **B:** SMPD4 expression is stimulated by Us9. Dissociated SCG neurons were infected with mock (uninfected), Us9^WT^, Us9^YY^, or Us9^null^ PRV strains to assay changes in SMPD4 protein expression. At 12hpi, samples were lysed and subject to SDS-PAGE for SMPD4 and Actin expression. SMPD4 expression is the strongest upon Us9^WT^ infection, followed by comparable levels in mock and Us9^YY^ infection, and lowest in Us9^null^ infection. **C:** SMPD4 localization in uninfected SCG neurons. Dissociated SCG cell body transduced with siRNA against Non-Target control (top) or SMPD4 (bottom) followed by immunofluorescence staining with DAPI, TGN38 for golgi and SMPD4.

